# Decoding P300 Variability using Convolutional Neural Networks

**DOI:** 10.1101/569616

**Authors:** Amelia J. Solon, Vernon J. Lawhern, Jonathan Touryan, Jonathan R. McDaniel, Anthony J. Ries, Stephen M. Gordon

## Abstract

Deep convolutional neural networks (CNN) have previously been shown to be useful tools for signal decoding and analysis in a variety of complex domains, such as image processing and speech recognition. By learning from large amounts of data, the representations encoded by these deep networks are often invariant to moderate changes in the underlying feature spaces. Recently, we proposed a CNN architecture that could be applied to electroencephalogram (EEG) decoding and analysis. In this article, we train our CNN model using data from prior experiments in order to later decode the P300 evoked response from an unseen, hold-out experiment. We analyze the CNN output as a function of the underlying variability in the P300 response and demonstrate that the CNN output is sensitive to the experiment-induced changes in the neural response. We then assess the utility of our approach as a means of improving the overall signal-to-noise ratio in the EEG record. Finally, we show an example of how CNN-based decoding can be applied to the analysis of complex data.

## 1 Introduction

Decades of neuroscience research have yielded profound insights into how the brain processes stimuli, integrates perceptual information, adapts to dual-task demands, and coordinates behavior. Using high-resolution electroencephalogram (EEG), researchers have examined the time course of multiple neural responses, including those related to stimulus encoding, cognitive processes, error detection, and motor movement. However, despite the extensive amount of research into such phenomena and the development of advanced imaging systems and signal processing techniques, the preferred method of analysis is still to collect large numbers of precisely timed trials within controlled experimental paradigms. This preference is due in part to the low signal-to-noise ratio (SNR) of task-relevant neural activity within the EEG record.

To improve the SNR of both oscillatory and evoked EEG signals, several neural decoding approaches have been developed. These approaches include methods such as Independent Component Analysis (ICA), Hierarchical Discriminant Component Analysis (HDCA), and Common Spatial Patterns (CSP), among others [1–4]. Whether these techniques are utilized for signal processing (e.g. ICA), or machine learning applications (e.g. CSP), they are calibrated using subject-specific training data. Such a requirement, however, reinforces the need for a large number of trials to reduce the likelihood of overfitting during the training process. Furthermore, these approaches make a number of assumptions about the data, such as non-Gaussianity, stationarity, whiteness, or statistically independent unitary dimensions, which may not extend to cross-session, cross-subject, or cross-experiment analyses. For example, ICA, CSP, and HDCA all utilize some form of spatial filtering, which can quickly become sub-optimal with slight variations in EEG electrode placement, as occurs across subjects or recording sessions [5]. ICA can converge on differing solutions for the same subject over different temporal epochs [6] or fail to capture signal variance due to enforcement of statistical independence [7]. In short, we are not aware of any neural decoding approaches that can be robustly applied across datasets or experimental paradigms, such that they do not require subject-, or at least experiment-, specific training data.

Recent advances in deep convolutional neural networks (CNNs) for EEG analysis [8–14] have opened the door for development and training of large-scale generalized models, i.e. models that can reliably decode neural activity across subjects and experimental configurations. In the broader machine learning community, CNNs have often been applied to pattern recognition problems such as automatic speech recognition and image processing (see [15–17] for reviews). CNNs make limited assumptions about the underlying data, can learn from large, diverse datasets, and extrapolate well to previously unseen data. Although generalized neural decoding models, such as those enabled by CNN architectures, have the potential to interpret data in more complex and less repeatable scenarios, as well as generally enhance SNR, it is important to quantitatively establish the link between the decoding model output and the underlying neural phenomena.

In this paper, we 1) establish such a link for the visual P300 evoked response, and 2) introduce a CNN-based approach for EEG decoding that can alleviate the need for large amounts of the test experiment’s data. We validate our CNN approach using a leave-one-out experimental analysis, and show that the outputs of the CNN (trained exclusively on other experimental datasets) faithfully replicate the well-known modulation of the P300 signal observed in the test set. The sources of P300 variability we examine include 1) perceptual similarity, 2) target-to-target interval, and 3) dual-task demands. We then show that the improved SNR in the CNN output space allows us to obtain the same level of significance when comparing experimental conditions using traditional statistical techniques, while requiring substantially fewer trials. Finally, we examine a free-viewing target detection task, in which the anticipated neural response is masked by both subsequent eye movements as well as task-relevant visual feedback. Using our CNN approach, the underlying target-related neural response can be recovered, whereas with traditional techniques, it cannot.

## 2 Review of the P300 Evoked Response

The P300 evoked response, first reported by Sutton in 1965 [18], is a stereotyped neural response to novel or task-relevant stimuli with maximal amplitude in the parietal region [19]. In a two-stimulus paradigm, the P300 is most often observed as a response to an infrequent target stimulus presented among more frequently occurring background stimuli with typically 1-3 seconds between stimulus presentations. In the three-stimulus version of the task, an additional infrequent ‘non-target’ stimulus is presented along with the target and background. In either case, only the target stimulus requires a response from the observer, though overt responses (e.g. button press) are not required to elicit the P300. When visual stimuli are used and the rate of stimulus presentations increases to 2 Hz or greater, the approach is commonly referred to as rapid serial visual presentation (RSVP) [20].

It is well established that both the amplitude and latency of the P300 are affected by endogenous, exogenous, and pathological factors [19, 21, 22]. This includes both feature (e.g. target and background similarity) and temporal (e.g. target frequency) stimulus properties, as well as cognitive states (e.g. high working memory load). As such, it provides an ideal event-related neural response from which to assess the effectiveness of our generalized neural decoding methodology. Prior research shows the P300 amplitude is generally larger with an earlier latency for tasks with easy versus difficult target/background discrimination, in tasks with long versus short target-to-target intervals, and in tasks with low versus high task demands.

### 2.1 Perceptual Similarity

Both the amplitude and latency of the P300 are modulated by the perceptual similarity between targets and non-targets. Specifically, P300 amplitudes in response to non-target stimuli tend to increase the more similar these stimuli are to target stimuli [23–26]. Variability in P300 responses evoked during an RSVP task can be attributed, in part, to the presence of distractor images that share physical and/or semantic characteristics with the rare target class of interest, interspersed with frequent background images [21]. Studies have shown that when the target/background discrimination difficulty increases, that is, when the target is similar to the standard background, P300 amplitude decreases and latency lengthens [27, 28].

### 2.2 Target-to-Target Interval

Prior work has also shown that P300 amplitude can change as a function of the target stimulus probability, the number of background images preceding the target (i.e. target-to-target interval or TTI), as well as the interstimulus interval (ISI) [29]. Targets at short ISIs tend to have smaller and longer latency P300 responses compared to those elicited at longer ISIs [30, 31]. However, when ISI is manipulated together with TTI, more of the P300 amplitude variance is accounted for by TTI than ISI [29]. P300 amplitude remains relatively constant when TTIs approach 6-8 seconds or longer [29]. Together these findings suggest that the neural system needs time to efficiently recover from processing target-related information.

### 2.3 Task Demands

Amplitude and latency effects have also been observed with changes in cognitive workload [32–34]. Specifically, it has been shown that working memory manipulations from one task can affect multiple levels of neural processing in another. For example, in a study by [33], subjects performed an arrow flanker task either alone or while performing a Sternberg task with high or low working memory load. The results showed decreased P300 amplitude with increased working memory load for the incongruent flanker stimuli. P300 latency is also affected by task demands, as demonstrated in a recent study by Ries et al. [35], which showed that target response latency significantly increased as a function of auditory working memory load.

## 3 Materials and Methods

### 3.1 P300 Database

Our P300 EEG database was constructed from four previously collected and analyzed P300 experimental datasets. All experiments were approved by the Institutional Review Board of the Army Research Laboratory. A high-level summary is given in Table 1. The database contains examples from both RSVP and free-viewing tasks, and were selected to investigate the decoding properties of our CNN approach. For our leave-one-out tests, by training a model using the three remaining datasets, we expected to sufficiently represent the variability in our hold-out set. An overview of the four experimental paradigms is shown in Figure 1.

**Table 1:**
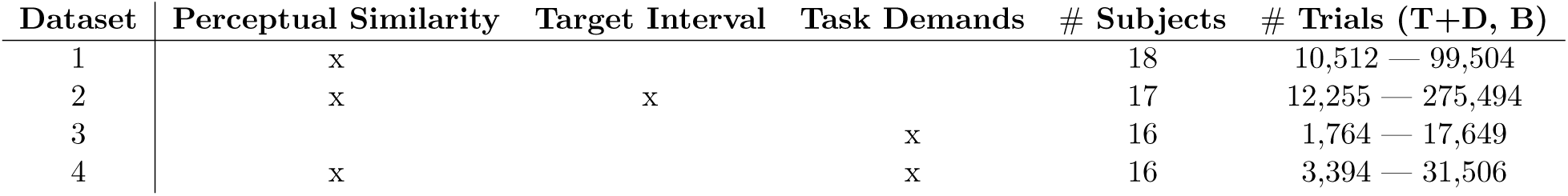
Sources of variability, # of Subjects, and # of Trials for each P300 Dataset (T = Target, D = Distractor, B = Background).

| Dataset | Perceptual Similarity | Target Interval | Task Demands | # Subjects | # Trials (T+D, B) |
| --- | --- | --- | --- | --- | --- |
| 1 | x |  |  | 18 | 10,512 — 99,504 |
| 2 | x | x |  | 17 | 12,255 — 275,494 |
| 3 |  |  | x | 16 | 1,764 — 17,649 |
| 4 | x |  | x | 16 | 3,394 — 31,506 |

**Figure 1:**
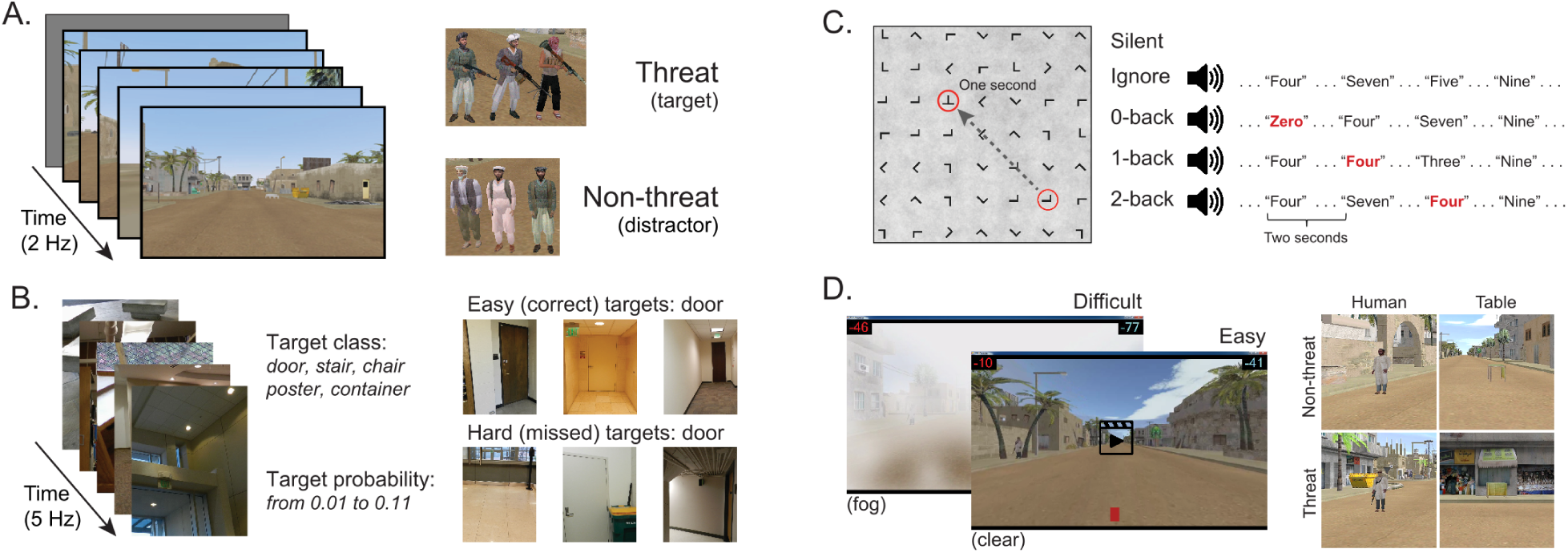
Experimental paradigms from the four datasets (A-D) used in this analysis.

#### 3.1.1 Dataset 1: Target Discrimination with Distractor Images

Our first dataset (Figure 1A) involved a 2 Hz RSVP task with targets, distractors, and background images [23]. Target to background ratio, and distractor to background ratio, were both approximately 1:14. Target images contained people holding a weapon, naturally positioned in a simulated urban environment, while distractor images contained people not holding weapons. Background images contained neither people nor weapons. To avoid interference from the attentional blink phenomena [36], at least two background images were required to follow any target or distractor image. Each subject experienced a total of four conditions for this dataset, which were the combinations of two manipulations: 1) subjects either mentally counted or pressed a button in response to targets, and 2) the run either had no distractors (only target and background images), or had all image types (target, background, and distractor images). In all conditions, the target as well as distractor stimuli could be stationary or moving. The appearance of moving stimuli was created by presenting five sequential images within the 0.5s stimulus epoch. In each image the target was in a slightly different location (i.e. animation). Data were collected from 18 subjects (13 male). EEG data were recorded with 64-channel BioSemi ActiveTwo (Amsterdam, Netherlands).

#### 3.1.2 Dataset 2: Target Discrimination with Complex Imagery and Variable Target-to-Target Intervals

Our second dataset (Figure 1B) is from a 5 Hz RSVP task that incorporated variations in TTI as well as stimulus complexity [37, 38]. Subjects performed six 10-minute blocks of target detection in which images of everyday office scenes were presented. There were five unique target categories: stairs, containers, posters, chairs, and doors. Before each block, the subject was notified of the target category for that block, and was instructed to push a button when a target stimulus was presented (i.e. go / no-go). There were six distinct experimental conditions where the TTI was varied. These six block types had average TTIs of 1.8, 2.2, 2.8, 3.9, 6.6, and 18.2 seconds. Targets also appeared at various sizes, eccentricities, and occlusion levels (i.e. target object could be fully visible or occluded by other objects in the scene), increasing the complexity of image categorization. Due to the difficulty of this task (on average, 44% of targets were missed), only trials with correct responses were used for this analysis. Data were collected from 17 subjects (7 male). EEG data were recorded with 256-channel BioSemi ActiveTwo. Channels were downsampled to the 64 BioSemi system montage.

#### 3.1.3 Dataset 3: Target Discrimination during Guided Visual Search under Varying Degrees of Cognitive Workload

The third dataset (Figure 1C) involved guided fixations around a grid of target and non-target stimuli, in both the presence and absence of an auditory task [35]. Eye movements were guided through the use of a red annulus across a grid of mostly L’s. The annulus moved to a new location approximately every second, and subjects made a button press when they fixated on the infrequent target letter, T. The guided visual search task was performed under five conditions: 1) alone (*Silent*), 2) while ignoring binaurally presented digits, numbers 0-9 (*Ignore*), or 3-5) while using the auditory digits in a 0, 1, or 2-back working memory task (*0-, 1-, 2-Back*). Auditory stimuli were presented every two seconds with a 500ms offset from the red annulus to prevent simultaneous auditory/visual events. Data were collected from 16 subjects (all male). Eye tracking data were acquired using a SMI RED 250 (Teltow, Germany). EEG data were recorded with a 64-channel BioSemi ActiveTwo. Horizontal and vertical electrooculogram (EOG) data were recorded, respectively, by placing electrodes near the outer canthus of each eye, and above and below the orbital fossa of the right eye.

#### 3.1.4 Dataset 4: Target Discrimination via Free-Viewing Visual Search during Video Playback

The fourth dataset (Figure 1D) is from a free-viewing task in which subjects viewed an urban landscape in a 15-minute video [39, 40]. In this video, subjects were driven through a simulated urban environment and required to find two different types of targets, humans and tables, and then discriminate between visually similar versions of each entity. Human entities were either holding a weapon (threat) or unarmed (non-threat). Tables were oriented in such a way that they could either hide an explosive device (threat) or not (non-threat). Target stimuli would abruptly appear one at a time in random but logical locations (i.e. on the street or in a doorway) at an approximate rate of once every three seconds, remaining on screen for one second. Subjects were free to scan the environment but were instructed to indicate the type of entity (threat or non-threat) by pressing a button with either the left or right index finger (i.e. two-alternative forced choice). Therefore, while the task was free-viewing, accurate discrimination tended to require that the subject fixate on the object. Participants were not required to hold target fixations for a prescribed amount of time, and thus subsequent fixations, along with other contaminating EOG effects, could occur once the subject had acquired enough information about the target to perform discrimination.

Subjects performed this task under two distinct visibility conditions: 1) a baseline of clear visibility (*Clear*) for easy stimulus detection, and 2) low visibility with an obscuring fog (*Fog*) for difficult stimulus detection. Subject responses were graded for speed and accuracy, and the subject received performance feedback for every target occurrence. The per trial score was visually displayed on the screen after button press response. Following the offset of each target stimulus, a cumulative score bar was updated, and displayed at the top and bottom of the screen. Data were collected from 16 subjects (all male). EEG data were recorded with a 64-channel BioSemi ActiveTwo. Horizontal and vertical EOG data were recorded in the same fashion as Dataset 3.

### 3.2 CNNs for Neural Decoding

CNNs for image processing typically require vast amount of training data, often on the order of millions of images distributed across thousands of classes [41–43]. In neuroimaging studies, however, collecting such vast amounts of data is prohibitively expensive and impractical, thus necessitating the need for CNN architectures designed specifically for low-sample neuroimaging data, while still being robust to both inter-and intra-subject differences. Recent work by [13] has yielded a CNN architecture for EEG (EEGNet) that can learn from relatively small amounts of data, on the order of hundreds of trials per subject. Furthermore, [13, 44] showed that EEGNet enabled cross-subject transfer performance equal to or better than conventional approaches for several EEG classification paradigms, both event-related and oscillatory. EEGNet is also the model used to obtain our cross-experiment results described in [26, 39, 45].

EEGNet is a compact CNN that takes minimally processed time-series EEG data as input, first using temporal convolutions that act as bandpass frequency filters, followed by depthwise spatial convolutions that act as spatial filters, which together improve SNR and reduce the dimensionality of the data. The depthwise convolution allows the model to learn spatial filters for each temporal filter, without being fully connected to all of the outputs of the previous layer and thus greatly reduces the number of parameters to be learned. EEGNet uses separable convolutions to more efficiently combine information across filters [46]. Each convolution layer is followed by batch normalization, 2D average pooling, and Dropout layers. We fit the EEGNet-4,2 model, denoting four temporal filters of length 64 samples and two spatial filters per temporal filter, as described in [13].

### 3.3 EEG Signal Preprocessing

For all EEG datasets analyzed in this paper, the data were bandpass filtered between 0.3 Hz and 50 Hz before being downsampled to 128 Hz. To ensure that all datasets were scaled similarly, we normalized each subject in each experiment by dividing the filtered data by its median absolute deviation (MAD), calculated over all channels and time points. We use MAD to reduce the effect of transient but large artifacts that occurred in some of the EEG recordings.

### 3.4 Model Training

All CNN models were trained using a leave-one-experiment-out procedure, combining data across all but one experiment for training and then testing on the held-out dataset. We used EEG epochs 1s in length for our training instances. For all datasets, with the exception of Dataset 4, instances were created by epoching [0,1]s around stimulus or fixation onset. In Dataset 4, the P300 responses appear to begin prior to fixation, and therefore training instances were epoched [-0.3 0.7]s around fixation onset [39].

Models were trained to perform a binary classification between instances that contained a P300 response (response to target (T) and distractor (D) stimuli), and and those that did not (responses to background stimuli (B)). When present, distractors were included in the target class to capture the natural variability of the P300 response, as they elicit attenuated P300 responses [24]. To handle imbalance in both class (more non-targets than targets) and experiment size (some experiments had more instances than others), we applied a sample weighting procedure during training to ensure each class and experiment contributed equally. This was done using the inverse proportion of the data in the training dataset. For example, if the target to background ratio was 1:4, the sample weight for all target trials was set to 4, while the sample weight for all background trials was set to 1. The weights to control for experiment imbalance were calculated in a similar manner. The weights to control for class imbalance and experiment imbalance were multiplied together to form the final sample weights used in model training. The EEGNet-4,2 model was implemented in Tensorflow [47], using the Keras API [46]. The model was trained for 100 iterations using the Adam optimizer with default parameter settings [48], with a minibatch size of 64 instances, optimizing a categorical cross-entropy loss function. The dropout probability was set to 0.5 for all layers. Source code for the models can be found at https://github.com/vlawhern/arl-eegmodels. We will refer to the EEGNet-4,2 model as the “CNN Model”, and its model outputs as “CNN Outputs”, for the remainder of the paper.

### 3.5 Model Testing

Once the model was trained, we applied it to the test set using the sliding window approach depicted in Figure 2. For all datasets, we made a prediction (using 1s of EEG as input) every six samples over a duration beginning at 1 second before, and ending at 1 second after, stimulus or fixation onset. For example, a CNN output at T=0 uses the EEG epoch [0,1]s as input, and a CNN output at T=0.5 uses the EEG epoch [0.5, 1.5]s as input. This produced a time-series of 65 CNN outputs, with each CNN output summarizing 1s of EEG data. Each output can be considered a probability that the *1s long input epoch of EEG data* contained the same neural response associated with target stimuli extracted from the training data. It is important to note that, depending on the step size, neighboring points may be highly correlated. For example, a step size of 100ms would produce neighboring points that were computed using data with 90% overlap. As a result, sharp deviations in the original signal, such as the relatively instantaneous phase resetting that initiates the P300 response, are difficult to pinpoint in time using our CNN outputs. Our modeling approach is more suited for assessing amplitude shifts that result from changes in the underlying signal as well as large latency shifts in the evoked response. Minor variances in latency, such as temporal jitter in the component waveforms, should be largely ignored by our convolutional approach. We show these sliding window outputs to provide a more complete perspective of model performance.

**Figure 2:**
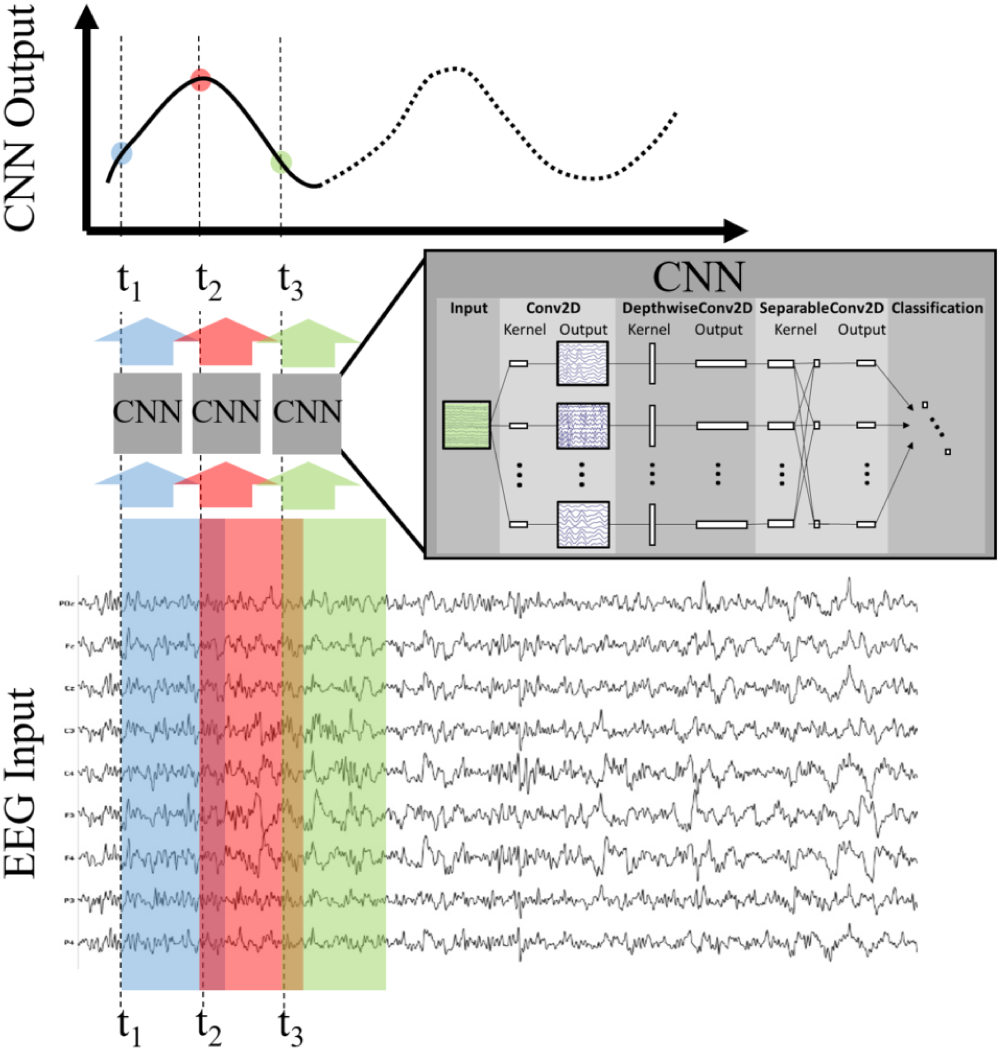
Production of CNN Outputs. Each CNN output is produced by passing as input an epoched sample of the EEG, i.e. a multi-dimensional array of channels × time. By stepping the CNN over the EEG record our approach produces an output signal that is decoded, in time and space, by the encoded deep model. For convenience purposes, we record the CNN output times using the *first* time point of the EEG epoch in question.

### 3.6 Statistical Testing for Signal to Noise Ratio Analysis

For our SNR analysis, using Dataset 3, we performed pairwise statistical testing on both P300 amplitudes as well as CNN outputs. The steps that we followed for testing P300 amplitudes were to 1) compute the average over a parietal region of interest (ROI) using channels Pz, P1, P2, CP1, CP2, and CPz in the 10-20 montage, 2) identify the mean value, per trial, in the window 400ms-850ms post-stimulus or fixation onset, 3) remove, per subject, the mean of (2) for background trials from the means computed for target trials, 4) aggregate the data across subjects, and then 5) compare the distribution of target means in the *Silent* condition to target means in the *2-Back* condition using a 1-tailed T-test. The steps that we followed to compare CNN outputs were similar to those used for P300 amplitudes, with the exception that we used the CNN output at *T* = 0*s* rather than means computed over time and space. In other words, we computed the average non-target CNN output, per subject, and subtracted that value from the per trial response for targets. We aggregated the target values across subjects and performed a T-test to compare the distributions of CNN outputs for targets in the *Silent* condition to targets in the *2-Back* condition.

In Section 4.4 we report the resulting p-values for Dataset 3 as a function of the percentage of trials selected. Trials were randomly downselected per subject, and only selected trials were used to perform both the P300 amplitude and CNN output tests. To reduce noise in our comparison resulting from trial selection, we repeated our random sampling and testing procedure 100 times for each percentage value analyzed.

## 4 Results

### 4.1 Perceptual Similarity

Figure 3A shows the P300 response, along with standard errors, as a difference wave for the target and distractor stimuli (i.e. target minus background, distractor minus background) for Dataset 1, aggregated across subjects. For this test, we compute our Evoked Responses and CNN outputs only for the condition that employs mental counting only (i.e. no button press) and has both target and distractor stimuli. The data for these plots was computed from the average over a parietal ROI (channels Pz, P1, P2, CP1, CP2, and CPz) and utilized the same preprocessing used to prepare the data for the CNN model. The CNN model was trained on Datasets 2, 3, and 4. Figure 3B shows the average CNN outputs for the same conditions presented in Figure 3A. We see that the model outputs closely mirror the P300 amplitude differences; in both plots, a clear separation of amplitudes is visible between target and distractor responses. In Figure 3B the CNN outputs peak at T=0, which is consistent with the model training, given that both use epochs [0,1]s around stimulus onset.

**Figure 3:**
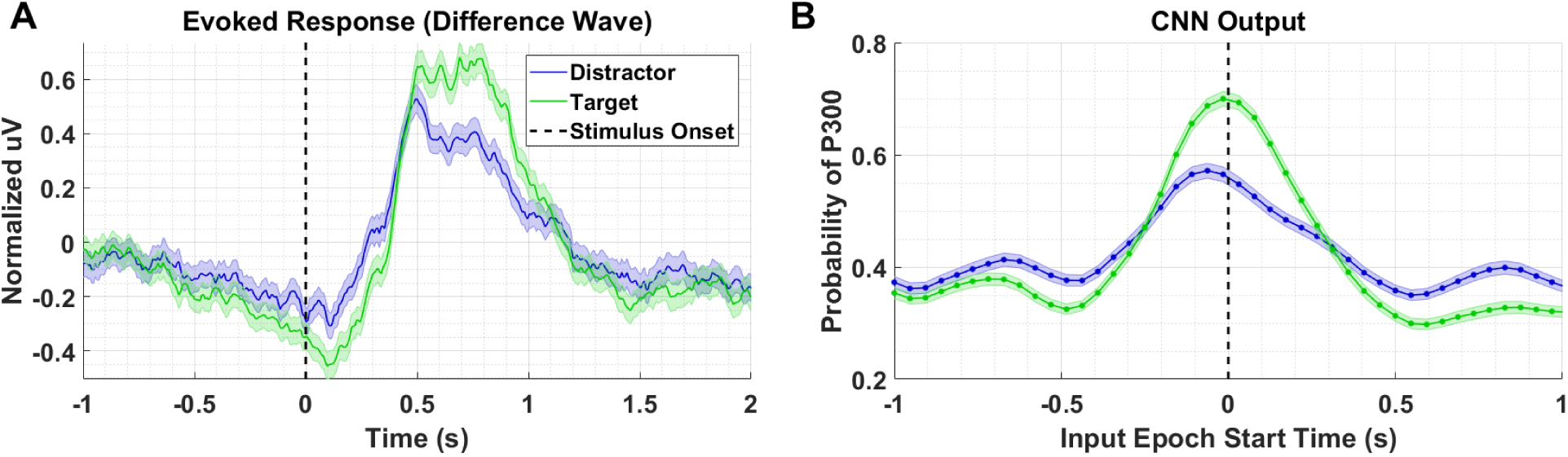
Perceptual Similarity Results using Dataset 1. A). Averaged P300 Evoked Response plots for target and distractors computed as difference waves for all subjects measured at the parietal ROI. B). CNN outputs are generated, for the same trials shown in (A), via sliding window approach detailed in Figure 2. Starting at 1s before stimulus onset, CNN outputs are generated (using 1s epochs of EEG data) every 6 samples, resulting in a time-series of 65 total predictions. Shaded regions in both figures denote 2 standard errors of the mean.

### 4.2 Target-to-Target Interval

Figure 4A shows the P300 response as a function of TTI for Dataset 2. Shorter TTIs (i.e. more frequent target presentations) measurably attenuate P300 amplitude. As TTI increases, the amplitude of the P300 response also increases. Figure 4B shows the model outputs for the same conditions presented in 4A. It should be noted that the responses in Figure 3A appear faster, yet persist longer, than the responses in Figure 4A. The baseline period, -0.5s to 0s, in Figure 3A shows greater desynchronization (i.e. negative dip). Although the origins of these differences are not critically important for our current analysis, it is important that the cross-experiment-trained models represented the amplitude shifts with a similar degree of fidelity, while also reflecting the subtle temporal shifts and overall shorter responses in Figure 4A. The CNN outputs in Figure 4B are shifted to the right and the main lobe appears thinner.

**Figure 4:**
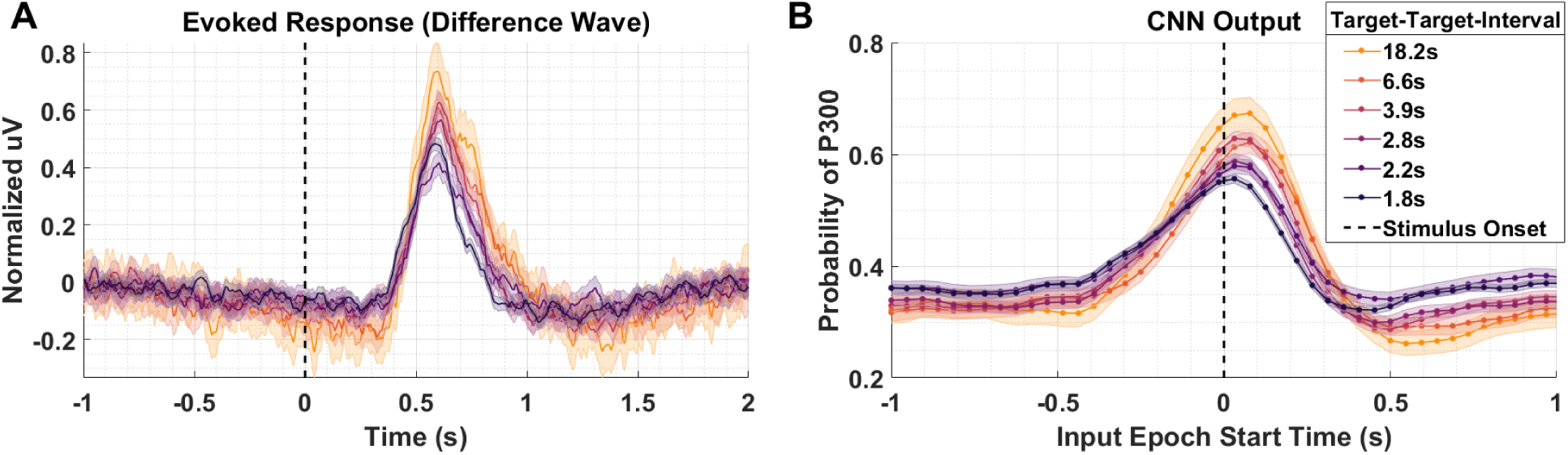
Target-to-Target Interval using Dataset 2. A). Averaged P300 Evoked Response (computed as difference waves) measured at the parietal ROI across subjects for the 6 distinct experimental conditions with varied TTI. The 6 conditions have, on average, TTI values of 1.8s, 2.2s, 2.8s, 3.9s, 6.6s, and 18.2s. B). A time-series of 65 CNN outputs are generated, for the same Target trials shown in (A), via sliding window approach detailed in Figure 2. Shaded regions in both figures denote 2 standard errors of the mean.

### 4.3 Task Demands

We compare the P300 waveforms from Dataset 3, in which subjects performed a guided visual search task while simultaneously performing an auditory N-back task at varying degrees of difficulty. Figure 5 presents the results for conditions *Silent, 0-Back*, and *2-Back* for both the P300 response (A) and model outputs (B). The high workload P300 waveforms have, on average, lower peak amplitudes and longer latencies. The model outputs reflect both of these shifts, though the shift in amplitude is much stronger than the shift in latency. That the model outputs continue to reflect shifts in amplitude and latency for fixation-locked P300 responses, when the majority of the training data were stimulus-locked, is further indication that the underlying template encoded within the CNN is based on a generalized representation of the P300 response. This template is reasonably invariant to the other experimentally-contingent components or induced artifacts, whether those are visual-evoked responses from rapid image presentations, or saccadic spikes/EOG contamination during visual search.

**Figure 5:**
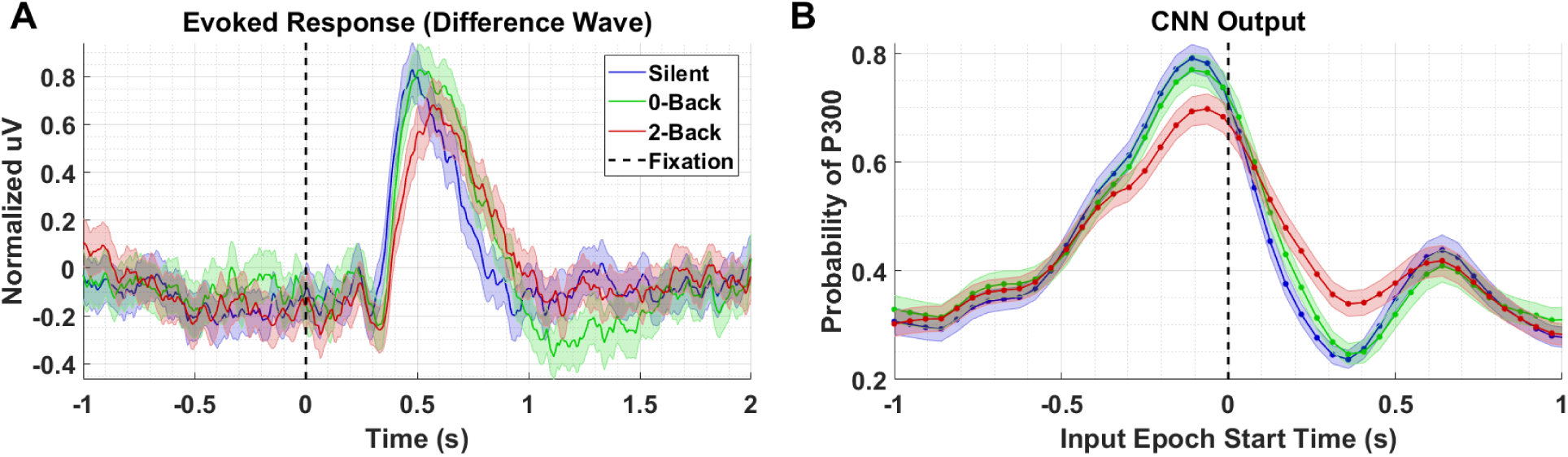
A). Averaged P300 Evoked Response (computed as difference waves) for targets trials, time-locked to fixation onset, obtained from the parietal ROI for the *Silent, 0-Back*, and *2-Back* conditions B). A time series of 65 CNN outputs are generated, for the same trials shown in (A), via sliding window approach detailed in Figure 2. Shaded regions in both figures denote 2 standard errors of the mean.

### 4.4 Noise Resistance

We performed the following analyses to assess the extent to which CNN-based decoding improves SNR. In the first test, we systematically downselected the amount of data extracted from a test set, Dataset 3, to compare the outcome of statistical testing as a function of the amount of data available. For this test, we compared data from the *Silent* and *2-Back* conditions; the results of this test are shown in Figure 6. As can be seen in the figure, the traditional analysis produces a p-value of approximately 0.01 using 100% of the data. An equivalent p-value can be obtained using the CNN output with roughly 25% of the data. Of course, the decision to reject the Null Hypothesis (i.e that target amplitudes are the same in both conditions) is a function of several factors, including significance threshold and multiple-comparison corrections. In the original paper [35], the authors did not report a significant difference in amplitudes after performing a multiple-comparisons correction. Use of CNN outputs would have affected this result due to the enhanced SNR of the CNN method.

**Figure 6:**
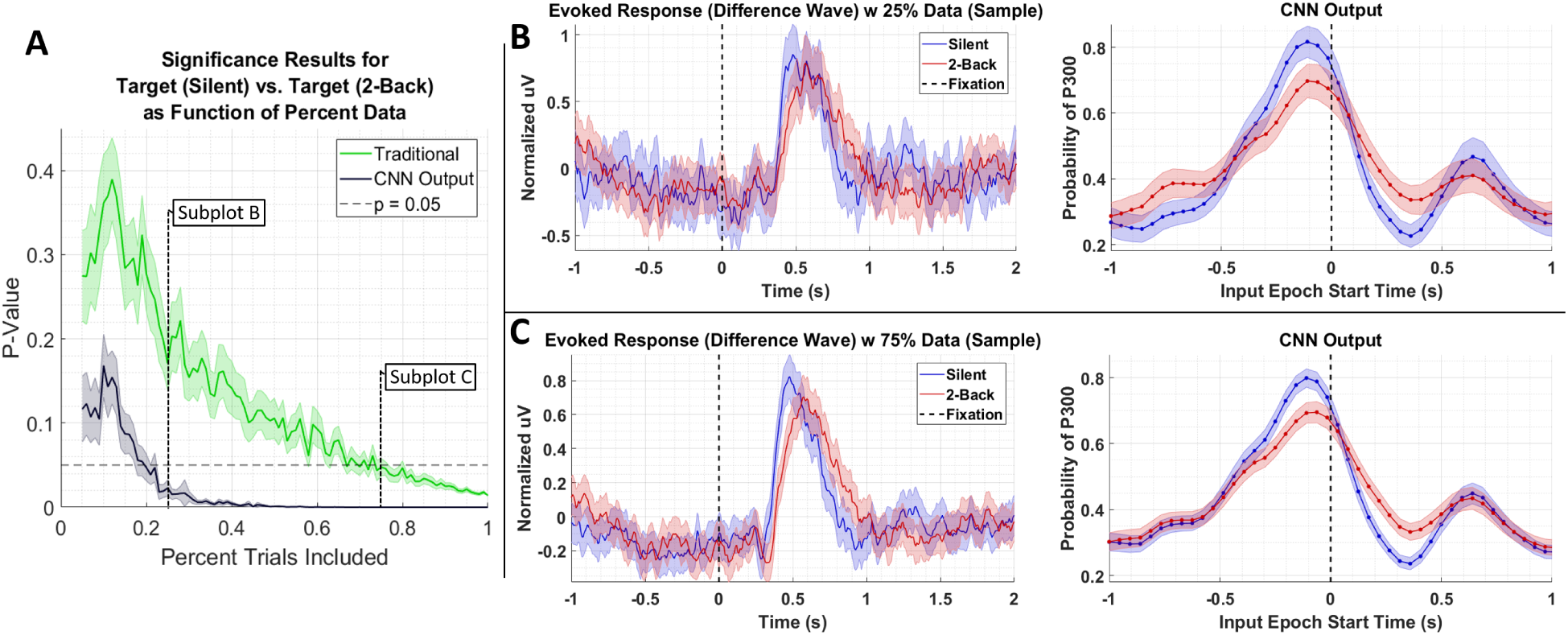
A). Result of significance testing for amplitude differences between target response in the *Silent* vs. *2-Back* conditions in Dataset 3 using CNN outputs (black) and measured P300 amplitude (green). B). Sample of the P300 and CNN outputs using 25% of the data. C. Sample of the P300 and CNN outputs using 75% of the data.

To better understand what is happening in the temporal domain, we performed a follow-up analysis on Dataset 2. We artificially created three groups of data from this dataset by grouping correct target responses by reaction time. The groups we considered were fast, medium, and slow RTs, which were calculated using the 33*^rd^* and 66*^th^* percentiles as cutoffs. These results are presented in Figure 7 and Table 2. As can be seen from Figure 7A, the P300 response for fast and medium responses appear relatively similar in amplitude, but the medium response is temporally shifted ∼100ms to the right of the fast response, which is consistent with the measured reaction times presented in Table 2. The P300 response for the slow group is shifted ∼100ms to the right of the medium response; however, the amplitude appears severely attenuated, while the lobe width appears wider. This phenomenon is largely a result of the unequal variance in the slow RT group; the P300 response in these trials is not necessarily diminished, but rather the increase in RT variance diminishes the amplitude of the averaged response [49]. Inspecting the results in Figure 7B, we observe that the time course of CNN outputs reflect the overall temporal trend of the three groups, i.e. the peak of the fast response is ∼100ms before the peak of the medium response, which is itself about ∼100ms before the peak of the slow response. In other words, a temporal shift of the entire distribution of the P300 response also shifts the CNN output peaks. Unlike the raw amplitude measurements, though, the CNN outputs indicate no difference in amplitude between the fast and slow responses. The convolutional structure of the CNN has, effectively, resolved the increased temporal variability in the slow group. There does appear to be a difference in amplitudes between the medium response and the response of the fast/slow groups; however, this can be explained by remembering that the CNN outputs are probability assessments that the input signal matches a pre-trained template, which is maximized in this case by the medium group. The shifts in amplitude as a function of latency are distributed about the center axis, i.e. T = 0s, in Figure 7.

**Table 2:**
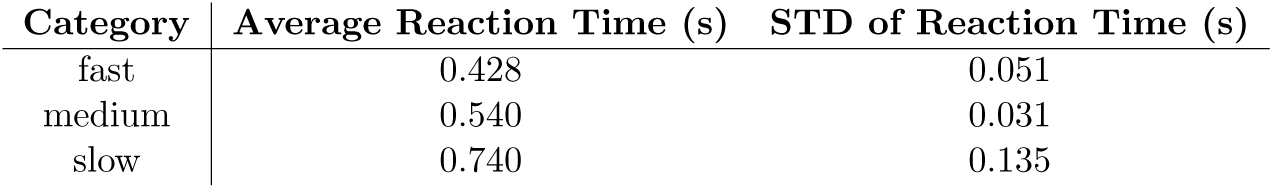
Mean and Standard Deviation values for Fast, Medium, and Slow reaction time groups.

**Figure 7:**
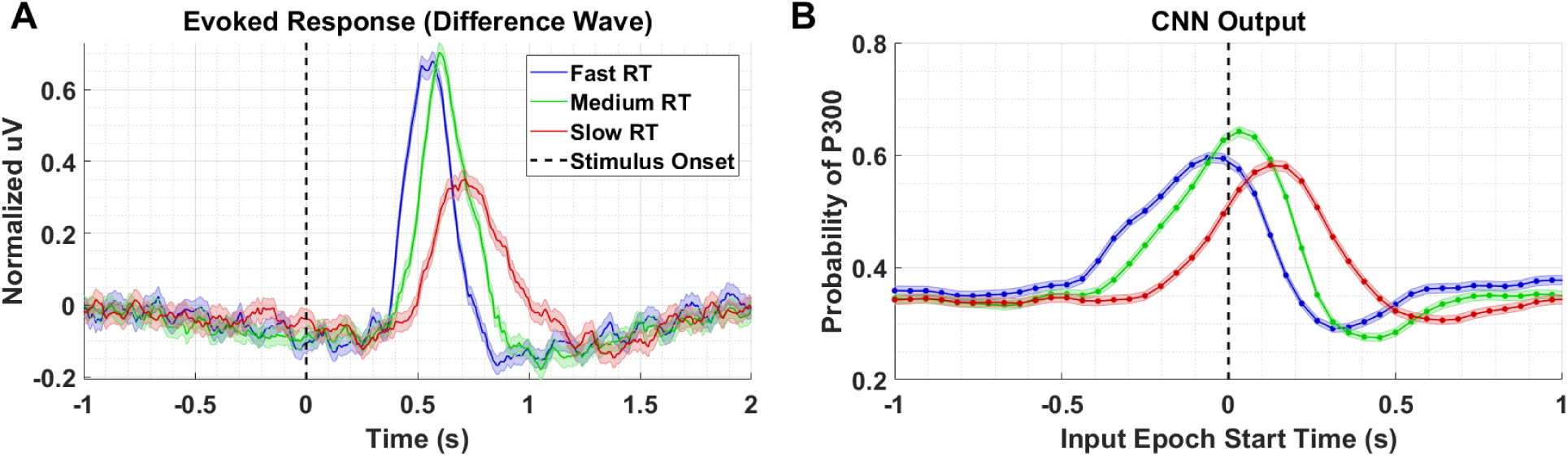
A). Averaged P300 Evoked Response (computed as difference waves) for correct target trials for each RT bin, detailed in Table 2 B). A time series of 65 CNN outputs are generated, for the same trials shown in (A), via sliding window approach detailed in Figure 2. Shaded regions in both figures denote 2 standard errors of the mean.

Returning to Figure 6B and C, we see that there is a minor, observable, shift in latency, which could impact the amplitude measurements of our CNN. However, the shift in latency for the low amplitude group (i.e. *2-Back*) is *towards* the center axis (T = 0s), where one would expect the CNN probability estimates to peak. In other words, the result in Figure 7 would suggest that if the amplitudes in Dataset 3 were equal, we would observe a lower amplitude for the *Silent* condition than the *2-Back* condition, due to the leftward shift of the *Silent* response. We argue that by observing the opposite trend, our confidence in the result increases.

### 4.5 Application to Complex Data

In our previous work [39] utilizing Dataset 4, we show that in the *Clear* condition, the fixation-locked and stimulus-locked neural responses are largely identical, with the exception of a small time shift. Here, the target’s “pop-up” effect acts an exogenous, or peripheral, cue that reliably guides the subject’s attention to the target. When the scene is obscured by fog, the latencies between stimulus onset and its subsequent fixation become longer and more variable, as the effect of this peripheral cue is diminished, and in some cases eliminated. As a result, subjects must employ more top-down search strategies to discover the target, which they may only perceive once it is within their parafoveal vision. Consequently, we expect that in the *Clear* condition, that the P300 response will be primarily locked to stimulus onset, whereas in the *Fog* condition, that the P300 responses are mixed between stimulus onset locking and fixation onset locking.

The latency between stimulus and a peripherally cued fixation is relatively stereotyped, with the most common fixation onset occurring 0.2194s post stimulus. We arrived at this value by fitting a lognormal probability distribution to a histogram of target fixation onset times in the *Clear* condition only, and choosing the peak value of the fitted distribution (the distribution of fixation onsets was heavily right skewed). In the *Fog* condition, we expect fixations that occur faster than this value to have predominantly fixation-locked neural responses, as these fixations occur faster than they would have if they were peripherally cued. In other words, considering the standard stimulus-to-fixation timing in the easier *Clear* condition, the subjects would not have had time to receive the peripheral cue and then saccade to the object. In Figure 8, we compare four conditions: *Fog* and *Clear* with short fixation onset times, and *Fog* and *Clear* with long fixation onset times. The *Fog* with short fixation times represents the case where the subject relies on no peripheral cuing. Short fixation onset times are defined as being less than 0.2194s, whereas long fixation onset times are defined as being greater than or equal to 0.2194s.

**Figure 8:**
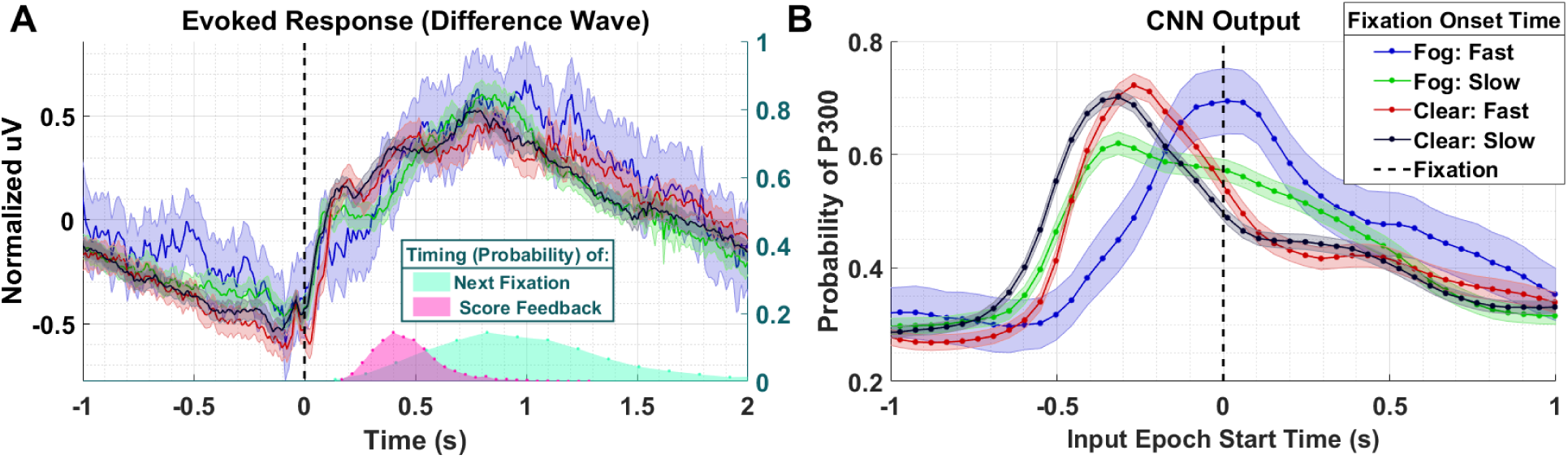
A). Averaged P300 Evoked Response (computed as difference waves) for target trials (left axis) time-locked to fixation onset, obtained from the parietal ROI and distribution of subsequent fixations as well as visual feedback time-locked to fixation onset (right axis). Fast fixation onset times represent those which occur faster than those which could have been peripherally cued B). A time series of 65 CNN outputs are generated, for the same trials shown in (A), via sliding window approach detailed in Figure 2. Shaded regions in both figures denote 2 standard errors of the mean.

In the Evoked Response difference waves shown in Figure 8A, the differences between the conditions are imperceptible. Importantly, the Evoked Response wave forms are atypical, with the apparent response persisting well after 1s post fixation, a result, we believe, of the mixture of subsequent search fixations (which occur ∼0.8s after target fixation) and visual feedback indicating the subject’s game score (∼0.5s after target fixation). In Figure 8B, we see that the CNN outputs for the *Clear* condition, both slow and fast, are similar to each other in timing and magnitude despite the difference in fixation latency relative to stimulus onset. However, the neural response, as measured by the CNN output, associated with short fixations (red-763 trials) occurs closer to fixation onset than the neural responses associated with long fixations (black-1,676 trials). For the *Fog* condition, we see that the CNN outputs associated short fixations (blue-101 trials) peak at T = 0s, which confirms our hypothesis that the neural responses are driven by top-down search rather than peripheral cuing. In other words, processing of these stimuli begin at fixation. For long fixation onsets in the *Fog* condition (green-854 trials) we see two peaks in the CNN output: one that begins prior to fixation onset, and one that begins at fixation onset. We hypothesize that the first of these peaks represents the *Fog* targets, despite their obfuscation, that subjects were able to detect through peripheral cuing. The second represents the trials with slow-fixation-onset where peripheral cuing did not play a significant role in target discovery.

## 5 Discussion

Traditional methods to analyze neural phenomena often require the collection of large numbers of precisely-timed trials due to the relatively poor SNR of minimally processed EEG data. Here we introduced a technique that utilizes convolutional deep-networks to decode EEG data. We applied this neural decoding approach to the analysis of the well-known P300 response. The P300 response has previously been shown to vary in both amplitude and latency as a result of a variety of experimental manipulations. We showed that the outputs of our CNN were sensitive to these changes, whether those changes occurred as a result of perceptual similarity, target interval, or task demands. We further showed that our CNN-based decoding approach improved overall SNR of the underlying EEG signal, allowing us to correctly reject the Null hypothesis for a comparison of amplitude means using only a fraction of the recorded data. We also demonstrated that the CNN outputs shift with latency of the underlying signal, but that temporal variability about a shifted mean does not impact CNN output to the same extent that it impacts averaged amplitudes. Perhaps most importantly, our technique can be completely pre-trained using previously run experiments. Taken together, these results validate the use of CNN architectures for generalized neural decoding and show that such decoding can substantially alleviate the need for large numbers of repeated trials.

However, there are limitations to our CNN decoding approach. Principally, for our chosen neural response, the P300 evoked response, the CNN model loses temporal resolution. This is a direct consequence of the windowing and temporal convolutions within the model that are designed to reduce the effects of temporal jitter. Whether this resolution is truly lost or must be further decoded by analyzing the hidden layers and internal activity of the network is an important question that should be addressed with future work. Of course, this also points at another potential drawback of the CNN-based approach, which is the overall interpretability of layered networks. Techniques for interpreting hidden layers and activity of deep networks have advanced greatly over the past few years, primarily in computer vision [50–53]. These techniques have recently been applied to neurophysiological data like EEG [13, 54], enabling additional insights into CNN-derived feature representations. However, algorithms and approaches to fully enable the interpretation and understanding of network behavior remains an open scientific question. In addition, while CNN decoding improves amplitude measurements in the presence of temporal jitter, large temporal shifts can create false changes in amplitude as a result of the template matching occurring within the CNN (see Figure 7 fast vs. medium response). The relationship between latency and measured amplitude can be a confound in P300 analysis. If the purpose of neural decoding includes amplitude comparisons at different latencies, one possible solution would be to tailor the training data to the anticipated latencies of interest.

Finally, as our decoding approach (1) can be pre-trained, (2) is capable of ignoring co-morbid, experimentally induced components or artifacts, and (3) exhibits a certain amount of invariance to temporal uncertainty, this work represents an important extension beyond the traditional approaches employed in the measurement and interpretation of neural phenomena. We believe the use of such CNN-based decoding will enable more complex, real-world neuroscience research [55–57]. By allowing the experimenter to develop decoding models from precisely controlled laboratory experiments, yet analyze data from a much smaller number of trials without requiring the same level of temporal precision, CNNs such as the one presented here, can help bridge the gap between our knowledge of how the brain functions in the laboratory and how it may function in the real-world. Although there remains additional work to develop, refine, and validate CNN decoding approaches, including different neural phenomena and brain states, we anticipate a growing reliance on deep models for neuroscientific research in complex, real-world environments.

## Acknowledgments

This project was sponsored by the US Army Research Laboratory under Cooperative Agreement Number W911NF-10-2-0022. The views and conclusions contained in this document are those of the authors and should not be interpreted as representing the official policies, either expressed or implied, of the US Government. The US Government is authorized to reproduce and distribute reprints for Government purposes notwithstanding any copyright notation herein.

